# Heterogenous organization in condensates of multiple transcription factors in embryonic stem cells

**DOI:** 10.1101/2024.06.14.599027

**Authors:** Azuki Mizutani, Cheng Tan, Yuji Sugita, Shoji Takada

## Abstract

Biomolecular condensates formed via liquid-liquid phase separation are ubiquitous in cells, especially in the nucleus. While condensates containing one or two kinds of biomolecules have been relatively well characterized, those with more heterogenous biomolecular components and interactions between biomolecules inside are largely unknown. This study used residue-resolution molecular dynamics simulations to investigate heterogeneous protein assemblies that include four master transcription factors in mammalian embryonic stem cells: Oct4, Sox2, Klf4, and Nanog. Simulations of the mixture systems showed highly heterogeneous and dynamic behaviors; the condensates mainly contained Sox2, Klf4, and Nanog, while Oct4 was dissolved into the dilute phase. Condensates consisted of loosely interacting clusters in which Klf4 was the most abundant in the cores. We suggest that Klf4 serves as a scaffold of the condensate where Sox2 and Nanog are bound to stabilize the condensate, whereas Oct4 is moderately recruited to the condensate, serving as a client mainly via its interaction with Sox2.

**Biological significance:** In eukaryotes, transcription is known to be regulated by many regulatory factors such as transcription factors, co-activators, and RNA polymerases, but precise molecular mechanisms of regulation remain obscured. A recently proposed model suggests that transcription-related proteins condense via liquid-liquid phase separation (LLPS) using their intrinsically disordered regions, which serves to control transcription. Master transcription factors in mammalian embryonic stem cells have been a model system. It was revealed that several master transcription factors exhibit LLPS by themselves, but dynamics and molecular structure of these proteins in their mixture have not been well addressed. In this study, we study molecular structures of condensates in a mixture of four master transcription factors, Oct4, Sox2, Klf4, and Nanog, via molecular dynamics simulation. We found that the three transcription factors Sox2, Klf4, and Nanog form mixed condensates, while Oct4 was largely dissolved. Klf4 mainly served as a scaffold of the condensate. The three proteins formed micelle-like structures as was recently found in the Nanog condensate. The condensates weakly recruited Oct4. Formation of heterogeneous condensates may provide fertile local environments in cells.

## INTRODUCTION

Biomolecular condensates formed via liquid-liquid phase separation (LLPS) of proteins and nucleic acids are ubiquitous in cells(1, 2). The condensate’s physicochemical nature and functional roles have recently been intensively studied. Many proteins in the condensates contain intrinsically disordered regions (IDRs), which are implicated to facilitate the LLPS. Aromatic, hydrophobic, and electrostatic interactions play crucial roles. In heterogeneous systems, some biopolymers can serve as scaffolds for the condensate, whereas other molecules may be recruited to the formed condensate, serving as clients. While condensates of systems containing one or two kinds of biomolecules have been relatively well studied(3, 4), those of more complex hetero-molecular systems are less characterized in depth at the moment.

In cells, a large fraction of molecular condensates is found in the nucleus, such as the nucleolus, Cajal body, and paraspeckle(1). In the context of transcription regulation, some transcription factors (TFs) are known to form droplets *in vitro* and are implicated to serve transcription regulation(5). In this transcriptional condensation model, gene regulation in mammalian embryonic stem cells (ESCs) has been a model system studied in depth by Young group(6); a small set of TFs, including Oct4, Sox2, Klf4, and Nanog, bound to super-enhancer elements and form condensates with coactivators, Mediators, and other proteins, regulating cell-type specific gene expression and thus controlling differentiation/pluripotency of ESCs(7).

However, due to the imaging resolution used in these studies, the details of molecular organization, their interactions, and dynamics within the condensates have not been well characterized. Molecular dynamics (MD) simulations can be complementary to imaging and other experiments in that MD simulations can visualize condensates at ultra-high spatiotemporal resolution(8-10). Especially for biomolecular condensates, a residue-resolution coarse-grained MD simulation is one of the promising approaches with their high efficiency and fair accuracy. Most commonly, in this context, each amino acid is represented by a single bead, and the interactions between amino acids are represented via bottom-up and physicochemical parameterization(11-15).

In our previous study, using the residue-resolution coarse-grained MD simulations, we investigated molecular mechanisms of condensate formation by Nanog(16), one of the core TFs in ESCs, which is known to form droplets *in vitro*(5). Nanog has two IDR domains flanking the DNA-binding domain. The C-terminal IDR contains repeats of pentapeptides (WR), including tryptophan residues (Fig. 1A). By simulations, we found that Nanog formed small clusters primarily via hydrophobic interactions within the WR regions and these clusters were bound to each other mainly via electrostatic interactions to form large condensate. In each cluster, the hydrophobic C-terminal IDRs tended to be located in the center of the cluster. In contrast, more hydrophilic DNA-binding domains and N-terminal IDRs were on the surface, making it similar to a micelle.

**Figure 1.**
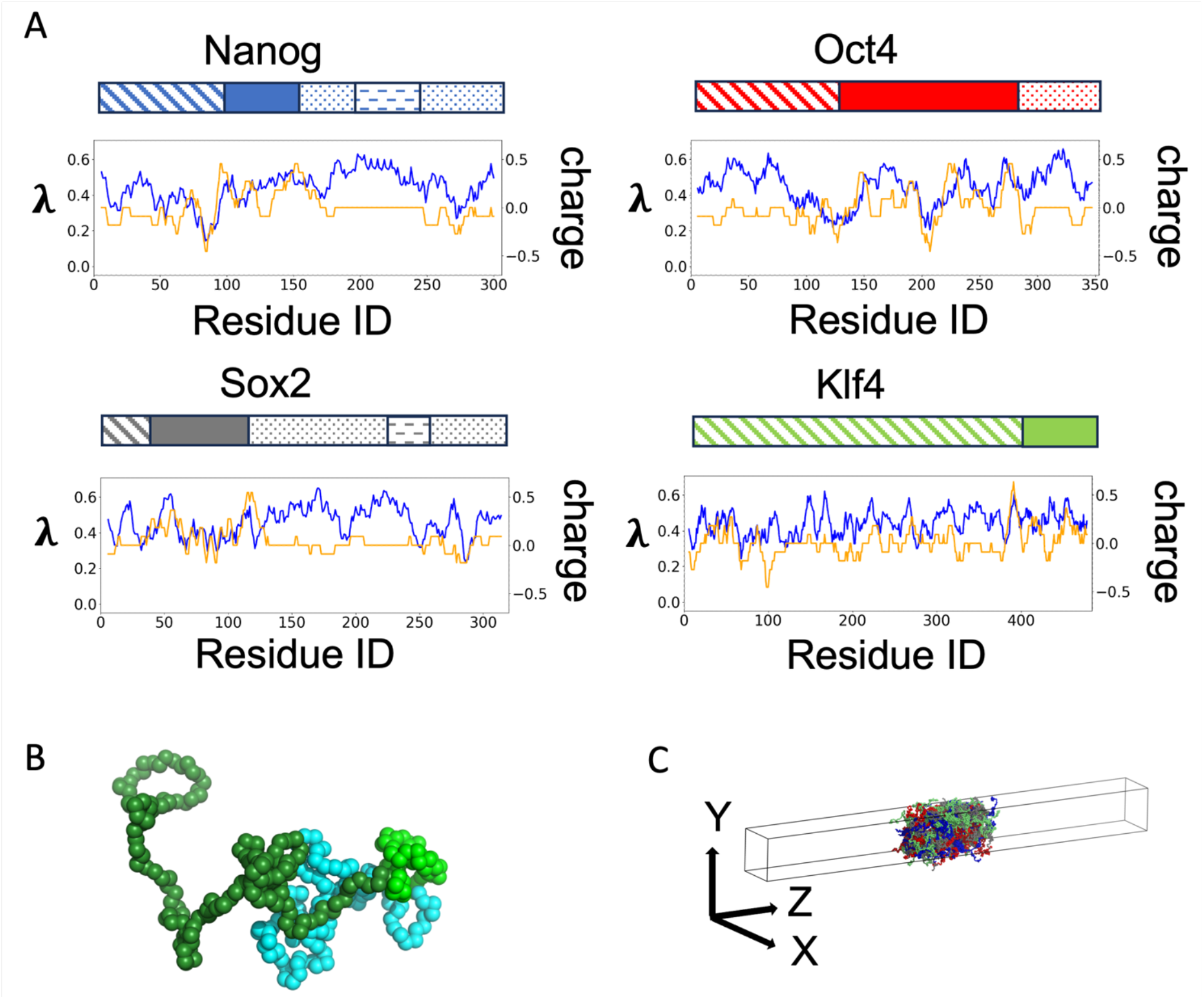
Four transcription factors, OSKN (Oct4, Sox2, Klf4, and Nanog), and simulation setup. (A) The domain organizations and one-dimensional hydrophobicity (blue) and charge (yellow) profiles for OSKN. The shaded, filled, and dotted rectangles represent N-terminal, DNA-binding, and C-terminal domains. The dashed region of Nanog and Sox2 represent a tryptophan-repeated (WR) region of Nanog and a tyrosine-repeated region of Sox2. For hydrophobicity (the HPS parameter) and charge, a moving average of 11 residues was plotted in the central residue position. (B) Residue-resolution model of Nanog. Each bead represents an amino acid. The cyan, green, and forest colored region represent the N terminal IDR, DNA-binding domain, and C-terminal IDR. (C) The slab simulation configuration.

In ESCs, several master TFs, including our current target proteins, Oct4, Sox2, Klf4, and Nanog (designated as OSKN), are highly expressed, making the clusters/condensates heterogeneous. How several molecules are organized in these cellular condensates are largely unknown. Among the four TFs OSKN, SKN are known to form droplets individually in *in-vitro* experiments; human Sox2, Klf4, and Nanog exhibited droplets at the average concentrations of 40, 2, and 10μM, respectively(5, 17). In the case of Nanog and Sox2, the polyethylene glycols (PEG) were used as a crowder at 10 % (weight percentage). On the other hand, human Oct4 showed no such droplets up to 40 μM(5). OSN showed some similarities in their domain organization (Fig. 1A). They all have a central DNA-binding domain flanked by two IDR domains, with the C-terminal IDR being generally more hydrophobic than the N-terminal IDR. Especially, the C-terminal domain of Nanog contains the WR region, which was suggested to form a core of micelles(16). Similarly, Sox2 has a tyrosine-repeat region in its C-terminal IDR. Unlike OSN, Klf4 possesses a long N-terminal IDR containing both charged and hydrophobic regions but no C-terminal IDR.

In this study, extending our previous work for Nanog(16), we investigated hetero-molecular protein mixture containing four master TFs of ESCs, OSKN. We address how the TF mixture is organized; do they form mixed condensates or separate condensates? In the mixture, what are the roles of each TF, scaffolds, or clients? What physical interactions are responsible for these behaviors? To address these questions, we began with modeling of each TF using a residue-resolution coarse-grained model (Fig. 1B). Then, we performed coarse-grained (CG) MD simulations for protein assemblies of individual TFs and the OSKN mixture in a slab-shaped configuration (Fig. 1C). We found that in the OSKN mixture simulations, Nanog, Sox2, and Klf4 formed the mixed condensates comprised of loosely interacting small clusters, while Oct4 were largely spread in the dilute phase. Comparison of the results of the OSKN mixture with those of individual systems suggested that Klf4 serves as a scaffold of the condensate, where Sox2 and Nanog are bound to stabilize the condensate, whereas Oct4 is moderately recruited to the condensate serving as a client mainly via its interaction with Sox2.

## RESULTS

### Sox2, Klf4, and Nanog, but not Oct4 form condensates in individual systems

Before investigating condensate formation in a mixture of OSKN, we first examined the propensity of each TF to form the condensates. For each mouse TF, we performed MD simulations for a system containing 200 molecules in a simulation box of 300 Å×300 Å×3000 Å at the 150 mM salt concentration, starting from completely phase-separated configurations (see METHOD for the initial configuration setup). We repeated five independent runs of 1.0×10^8^ MD steps with different stochastic forces.

In Fig.2A∼D, the top panels depict the time courses of the center of mass of every molecule along the long axis (Z-axis) in one trajectory (The results for other trajectories are shown in Fig. S1, S2). We found that Sox2, Klf4, and Nanog largely kept the condensates, while Oct4 spread uniformly, which is consistent with the previous experiments(5, 17). The time courses of the number of molecules in the condensates suggested that the trajectories reached near-equilibrium after ∼5.0×10^7^ MD steps for each TF (Fig.2 A∼D, middle panels, red curves). In the case of Klf4, the mean value of the number of molecules in the condensates was 184±3, while those of Nanog and Sox2 were lower, 153±4 and 113±8, respectively. Thus, Klf4 had a higher propensity to form the condensate than Sox2 and Nanog. This is consistent with the experimental results that Klf4 formed droplets with a lower concentration than Sox2 and Nanog(17). Klf4 has a longer IDR with many hydrophobic and charged segments than Sox2 and Nanog, which may stabilize the condensates of Klf4. Notably, Nanog in the current simulation with a salt concentration of 150mM showed a condensation propensity slightly, but non-negligibly, weaker than that we found in the previous work(16) that used the salt concentration 125mM motivated from the *in vitro* experiment (Fig. S3). The snapshots of the final frame showed that molecules did not form a single condensate but split into several condensates (Fig. 2A-D, bottom images). To study the detail of the condensates, we plotted the time courses of the maximum cluster size in the same figure (Fig. 2A-D, middle panels, blue curves). In the case of Klf4, the maximum cluster size was close to the number of molecules in the condensates, while those of Sox2 and Nanog were much lower than the size of the condensates. The longer IDR of Klf4 can interact with more proteins, which makes it easier to stabilize larger condensates. On the other hand, Sox2 and Nanog tend to split into smaller clusters at this condition.

**Figure 2.**
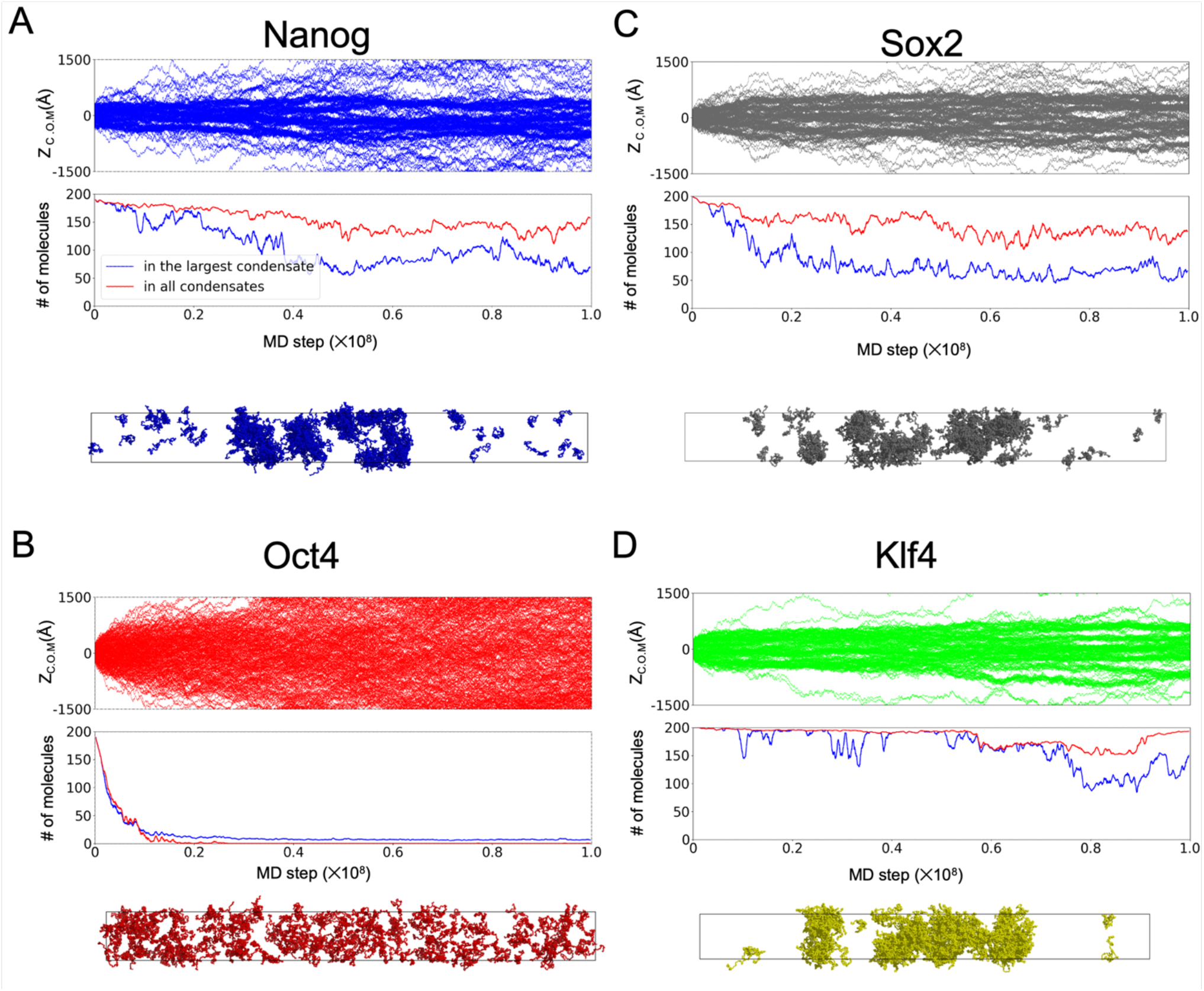
Simulations of individual TF systems containing 200 molecules. In each of A∼D, results of representative trajectories were plotted. (top) The time courses of Z-coordinates of the center of mass of individual molecules. (middle) Numbers of molecules in all the clusters (red) and those in the largest cluster (blue), plotted using a moving average of every 51 frames. (bottom) The snapshot at the end of the simulations. A; Nanog, B; Oct4, C; Sox2, D; Klf4.

### Sox2, Klf4, and Nanog mixture form heterogenous condensates

Next, we investigated the mixture of four TFs, OSKN. We set up the system that contains 50 molecules of each TF in the same slab simulation configuration as above. As a preparation, we first put all proteins in a box of size of 300 Å×300 Å×5000 Å and performed a short simulation shrinking the box size to obtain a compact mixture, which was followed by long relaxation simulations (1.0×10^8^ MD steps, five independent runs) with a just-sized box (see METHOD for the detail of the optimization). Using the final structure in the preparation as the initial configuration (Fig.3C top), we conducted production runs for 1.0×10^8^ MD steps within the 300 Å×300 Å×3000 Å box five times with different stochastic forces.

In the product runs of the four TFs, SKN molecules kept high densities around the center of the simulation box, while Oct4 rapidly spread nearly uniformly (Fig. 3A, Fig. S4A∼D). Of the SKN, Klf4 tends to be the most condensed. Sox2 and Nanog showed weaker propensities to stay in the condensate. Notably, during the simulations, small number of Oct4 molecules stayed in the condensates, suggesting that Oct4 is not completely excluded from the condensate. The number of molecules in condensates dropped from its initial value of 200 to ∼130±5 because most of the Oct4 molecules and a small fraction of SKN diffused to the dilute phase (Fig. 3B, Fig. S4E). The number of proteins in the condensates fluctuated between 120 and 150 after 6.0 ×10^7^ step, reaching nearly converged states. However, the largest cluster size gradually decreased until the 1.0×10^8^ MD steps simulation ended. Thus, we extended a simulation to 2.0×10^8^ steps, in which the largest cluster size reached near equilibrium (Fig. 3B inserted panel). In the extended trajectories, the largest cluster size was around 100, and the number of molecules in the condensate phase was 120 ∼ 150, which suggested that the condensed phase was not a single cluster. Still, each cluster contained tens to a hundred molecules.

**Figure 3.**
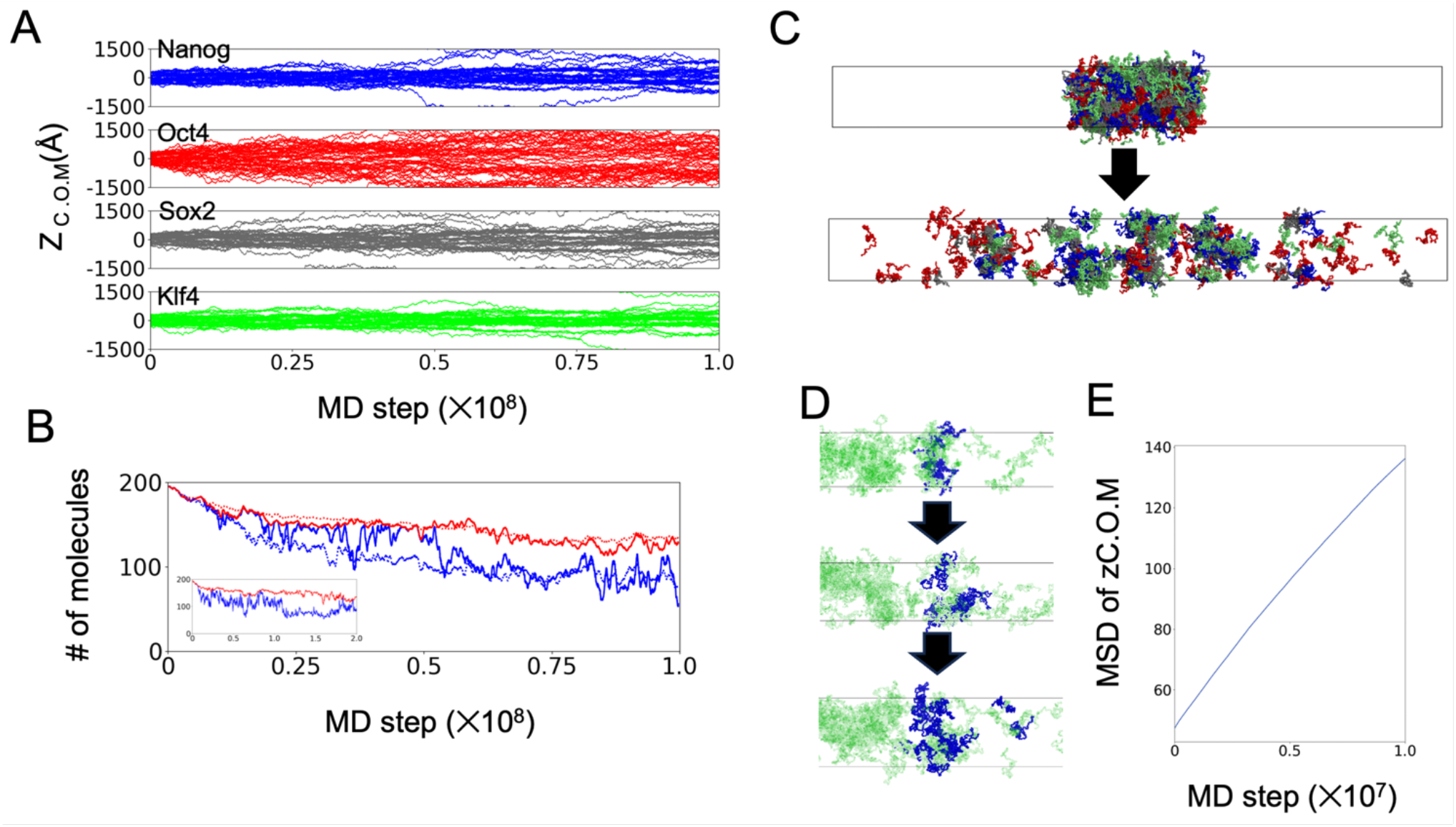
Simulations of OSKN mixture containing 50 molecules of each TF. (A) The time courses of Z-coordinates of the center of mass of individual molecules. Each panel shows the result of Nanog, Oct4, Sox2, and Klf4 from the top to the bottom. (B) The moving-averaged numbers of molecules in the clusters (red) and those in the largest cluster (blue). The dashed curves represent the average over five trajectories. Inset: Results of a doubly extended run. (C) The snapshots of the first and final structures in the same trajectory as those in Fig. 3A. In each snapshot, Nanog, Oct4, Sox2, and Klf4 were colored blue, red, gray, and green. (D) Tracking molecules in the same cluster at the top panel (blue molecules). The middle and bottom panels are the snapshots after 0.5×10^7^ and 1.0×10^7^ MD steps from the top panel. (E) The growth of the mean squared deviations of the Z-coordinates of the COM of the molecules in the same bin at the start point averaged over trajectories.

Visual inspection of snapshots (Fig.3C) suggested that condensates containing OSKN split into several loosely and dynamically interacting clusters. SKN molecules were mixed up to form the clusters rather than forming separate clusters of individual TFs (Fig. 3C). Previous studies showed that SKN proteins were colocalized at the target gene loci in chromatin(18). Our results suggest that SKN proteins can mix to form condensates even without a chromatin environment.

During the product runs, all the proteins moved dynamically with marked diffusion. Proteins in the dilute phase frequently merged into condensate, and proteins in the condensate often went out (Fig.3A, SI movie). To evaluate the liquidity of the condensates, we examined the diffusional movement in the condensates using a FRAP-like analysis(19, 20). At a frame, we color-coded molecules that were in a certain bin along the z-axis (Fig. 3D top panel). We tracked the motions of these marked molecules in subsequent time (Fig. 3D bottom panel). The standard deviation of the Z-coordinate center of mass of the marked molecules increased linearly in time (Figure 3E), which indicated the normal diffusion and that the condensates were liquid-like, made via the liquid-liquid phase separation.

### Heterogeneity in cluster cores

During the simulations of the OSKN mixture, SKN proteins did not maintain one large condensate but split into several clusters. We observed occasional splits and coalescences of clusters, reaching a nearly steady state by the end of simulations. To address the robust and key features of the clusters, we tried to exclude the fragile interactions and focused on the interaction in the “cluster core”, which was defined by a more stringent criterion of protein-protein interactions than the criterion of a regular “cluster” (see “Analysis” for details).

We first examined fractions of OSKN in the cluster core (Fig. 4A). While most SKN molecules were in clusters with loose interactions, their fractions in the cluster core were different among SKN. The most abundant TF was Klf4, comprising ∼40%, followed by Sox2 and Nanog. Oct4 constitutes only for ∼5% of all proteins located in the core. Notably, the content of Sox2 was higher than that of Nanog, which is opposite to the order of the propensity to form the condensate in the individual systems (Fig. 2), implying the importance of interactions with other kinds of TFs, especially with Klf4.

**Figure 4.**
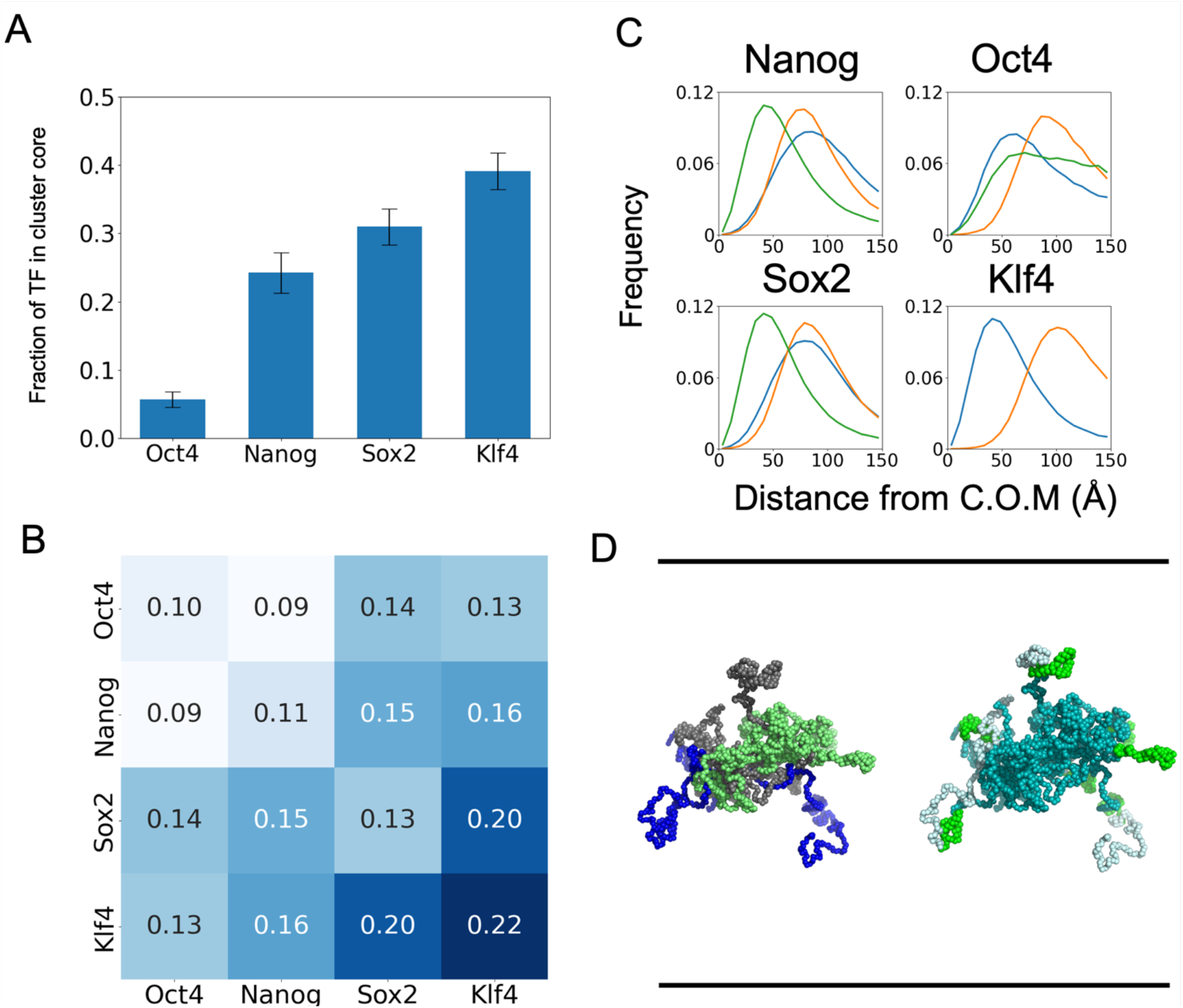
Heterogeneous organization of OSKN in the cluster core. (A) The fraction of TFs in cluster cores. The error bars are standard errors based on the results of the final frame of five trajectories. (B) The TF-TF interaction propensities. (C) The distribution of the distances of TF domains from the COM of cluster cores in the cluster cores. The blue, yellow, and green curves represent the results of N-terminal IDRs, DNA-binding domains, and C-terminal IDRs, respectively. (D) Representative snapshots of a typical single cluster core. In the top panel, residues were colored by molecules: Blue; Nanog, gray; Sox2, green; Klf4. In the bottom panel, the residues were colored by domain types: Cyan; hydrophobic IDRs, green; DNA-binding domains, gray; relatively hydrophobic IDRs. All the results used data from 5 trajectories. Horizontal bars represent the box size in X and Y axes, 300 Å.

To examine the detail of the interactions between the same and different TF molecules within the cluster core, we defined the contact propensity between each pair of TFs (Fig. 4B). Based on the intermolecular contact map used to define the “cluster core”, we counted the number of contacts between each pair of TFs relative to the number of all molecular pairs of the same TF pairs within the cluster core. Results showed the highest contact propensity being the Klf4-Klf4 contact. Klf4 has high contact propensities to Sox2 and Nanog as well, suggesting that Klf4 served as a scaffold of the mixed condensate. We noticed that the A-B contact propensities generally tend to be between those of A-A and B-B, with some exceptions (here, A and B stand for TFs). The contact propensity of Sox2 and Nanog was higher than those of their self-contacts, i.e., Sox2-Sxo2, and Nanog-Nanog. Sox2 is known to attract Nanog by the aromatic/hydrophobic interactions between the C-terminal IDRs of both TFs(21). While Sox2 and Nanog can induce LLPS by themselves, their cross-interaction seems stronger than the self-interaction. Similarly, Oct4 has a low contact propensity to Oct4 itself (0.10) but has a higher value with Sox2 (0.14), suggesting a specific pairwise interaction between Oc4 and Sox2. Notably, Oct4 and Sox2 are known to form hetero-trimers with DNA, with positive cooperativities(22). The current results suggest that, even without DNA, they have a weak attraction. Since Sox2 can form the condensate, the interaction between Sox2 and Oct4 may be a driving force for Oct4 to be recruited to the condensate.

So far, we have characterized the contact propensity at the resolution of molecules. Next, we investigated the localization of every domain/region within the “cluster cores”. In the analysis, we divided each TF into the N-terminal IDR domain (NTD), the globular DNA-binding domain, and the C-terminal IDR domain (CTD) (Klf4 has no CTD). We calculated the distance of each domain from the center of mass of the “cluster core” (Fig. 4C). We found that CTDs of Nanog and Sox2 were closer to the center, while NTDs were on the surface of each “cluster core”. In the case of Klf4, the N-terminal IDR is in the center of the “cluster core”, and its DNA-binding domain is on the surface of the “cluster core”. Comparing the properties of IDRs for each TF, we found that the average hydrophobicity of the IDRs was different. As shown in Fig. 1A, CTDs of Nanog and Sox2 and NTD of Klf4 have higher hydrophobicity, while NTDs of Nanog and Sox2 have lower hydrophobicity and more charges. Thus, different physicochemical properties of NTD and CTD of Nanog and Sox2 resulted in different localizations in the “cluster cores”. Relatively hydrophobic IDRs interact with each other to form “cluster cores” and hydrophilic IDRs and DNA-binding domains are on the surface of the “cluster cores”, suggesting a micelle-like architecture previously found in the Nanog condensate(16). A representative snapshot of the cluster core (Fig. 4D) illustrates that hydrophobic IDRs (dark green regions) interact with each other in the “cluster core”, while DNA-binding domains (green region) and hydrophilic IDRs (white region) are on the surface of the “cluster core”. Our previous study found that Nanog alone formed similar clusters by the hydrophobic interactions between the tryptophan-repeated (WR) regions in the CTD IDRs(16). Similarly, the features of “cluster cores” formed by SKN are consistent with the typical physiochemical nature of micelles. Our previous studies showed that the micelle-like clusters formed by Nanog could make DNA fragments closer(16). We propose that the formation of micelle-like clusters by SKN is important for the enhancer-promoter interaction.

### Residue-specific interactions in the cluster core

We have seen that OSKN have distinct preferences for the condensate formation, and each TF has distinct domains responsible for the condensate formation. Here, we further scrutinized the interaction at the resolution of individual residues. We analyzed reside-wise contact frequencies (Figure 5). We show residue-pairwise contact frequencies within the cluster cores in Fig. 5A∼C and Fig. S5. We also plotted the average contact probabilities of every residue (Fig.5D). We used the last 5000 frames of all five trajectories for these analyses. We removed molecules in the dilute phase from the analysis.

**Figure 5.**
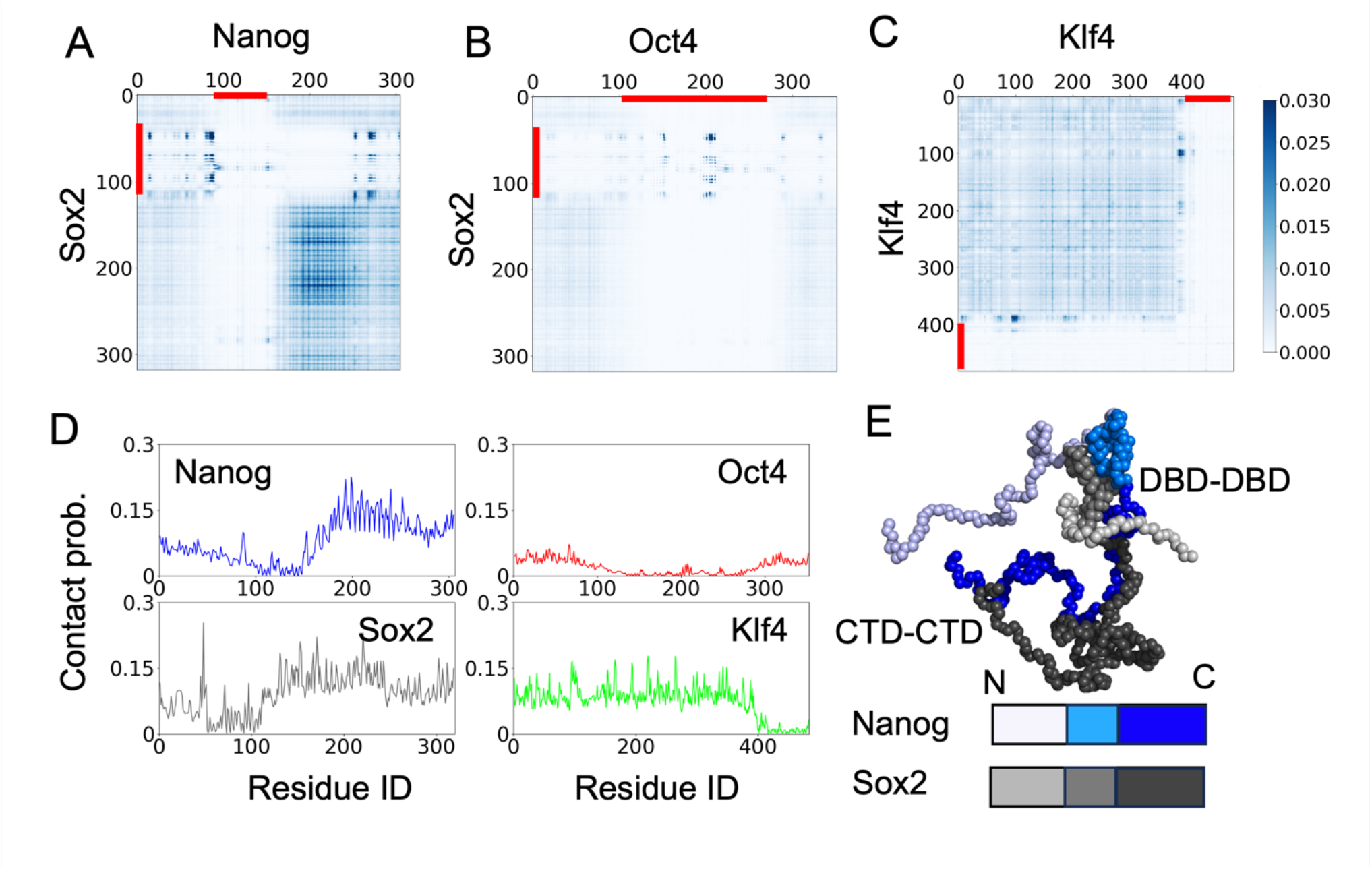
Residue-wise TF-TF interactions. (A∼C) The average number of contacts between residues of each pair of TFs. The red bars represent the DNA-binding domains. (D) The contact probability of each residue for each TF. (E) A snapshot of a complex of Nanog and Sox2. The colors are divided into the N-terminal tail, the DNA-binding domain, and the C-terminal tail. All the results in A∼D used data from five trajectories.

For the Sox2-Nanog interactions (Fig. 5A), frequent contacts were found between CTD of Sox2 (120∼230 residues) and CTD of Nanog (155∼240 residues). Both CTDs have high average hydrophobicity and few charged residues (Fig. 1). Especially the 195∼245 residues of Nanog and 221∼234 residues of Sox2 are known as the tryptophan-repeated (WR) regions and the tyrosine-repeated regions, respectively, containing repeated aromatic residues. These regions have high average contact probabilities (Fig.5D). Previous studies showed that Nanog and Sox2 formed heterodimers through interactions between the two regions(21). Our results are consistent with this result. We illustrated one snapshot of Sox2-Nanog dimer within the cluster core in Fig. 5E. In the snapshot, Nanog interacts with Sox2 in two separate regions. In one, IDR CTDs of Nanog and Sox2 interact with each other mainly via aromatic residues. In addition, we found contacts between two DNA-binding domains. Therein, Sox2 has some negatively charged regions, which interact with the positively charged region next to the DNA-binding domain of Nanog (Fig. 5A).

Figure 5B shows the interactions between Sox2 and Oct4. The contact frequencies were low overall because there were few Oct4 molecules in the cluster core. Yet, we found marked contact frequencies in the DNA-binding domains: Sox2 (40 - 112 residues) and Oct4 (131 - 280 residues). DNA-binding domains of both TFs have positively charged surfaces for binding to DNA. In addition, Oct4 has negatively charged residues around residue 200. The electrostatic interaction between the negatively charged residues of Oct4 and the DNA-binding domains of Sox2 contributed to the specific pairwise interaction of Oct4 and Sox2.

Compared to Sox2 or Nanog, we did not find specific regions important in the interactions between Klf4 molecules (Fig. 5CD). In the IDR of Klf4, both hydrophobic residues and charged residues are widely distributed (Fig. 1). We suggest that the tendency of Klf4 to induce LLPS is stronger compared with other TFs because Klf4 can interact with each other across long and multifaceted IDRs (Fig.5D).

### Electrostatic interaction in the condensate

In the previous section, we highlighted the role of hydrophobic and the electrostatic interactions. Here, we investigated the role of electrostatic interaction by conducting the simulations at different salt concentrations. A higher salt concentration screens the Coulomb interactions, resulting in the weakening of electrostatic interactions. We performed simulations of the same setup as above with different salt concentrations: 100mM and 500mM NaCl concentrations. Note that the above results were obtained with 150mM.

Figure 6A shows representative time courses of the number of molecules in clusters for three salt concentrations. Clearly, the number of molecules in the clusters decreases with the salt concentration. Thus, electrostatic interactions significantly contributed to the cluster/condensate formation. Figure 6B shows the snapshots of structures at the end of simulations with NaCl concentrations of 100mM and 500mM. With 500mM NaCl, clusters became smaller, and many molecules were in the dilute phase. In our previous study, clusters of Nanog were also observed in the case of 500 mM NaCl. It was because Nanog formed small clusters through hydrophobic interactions between the WR regions. This system also contained 50 Nanog molecules, and as shown in the previous sections, the “cluster cores” were thought to be formed primarily via the hydrophobic interactions between the relatively hydrophobic IDRs. We hypothesized that the condensates are broken by increasing the salt concentration, but the cluster cores might remain intact.

**Figure 6.**
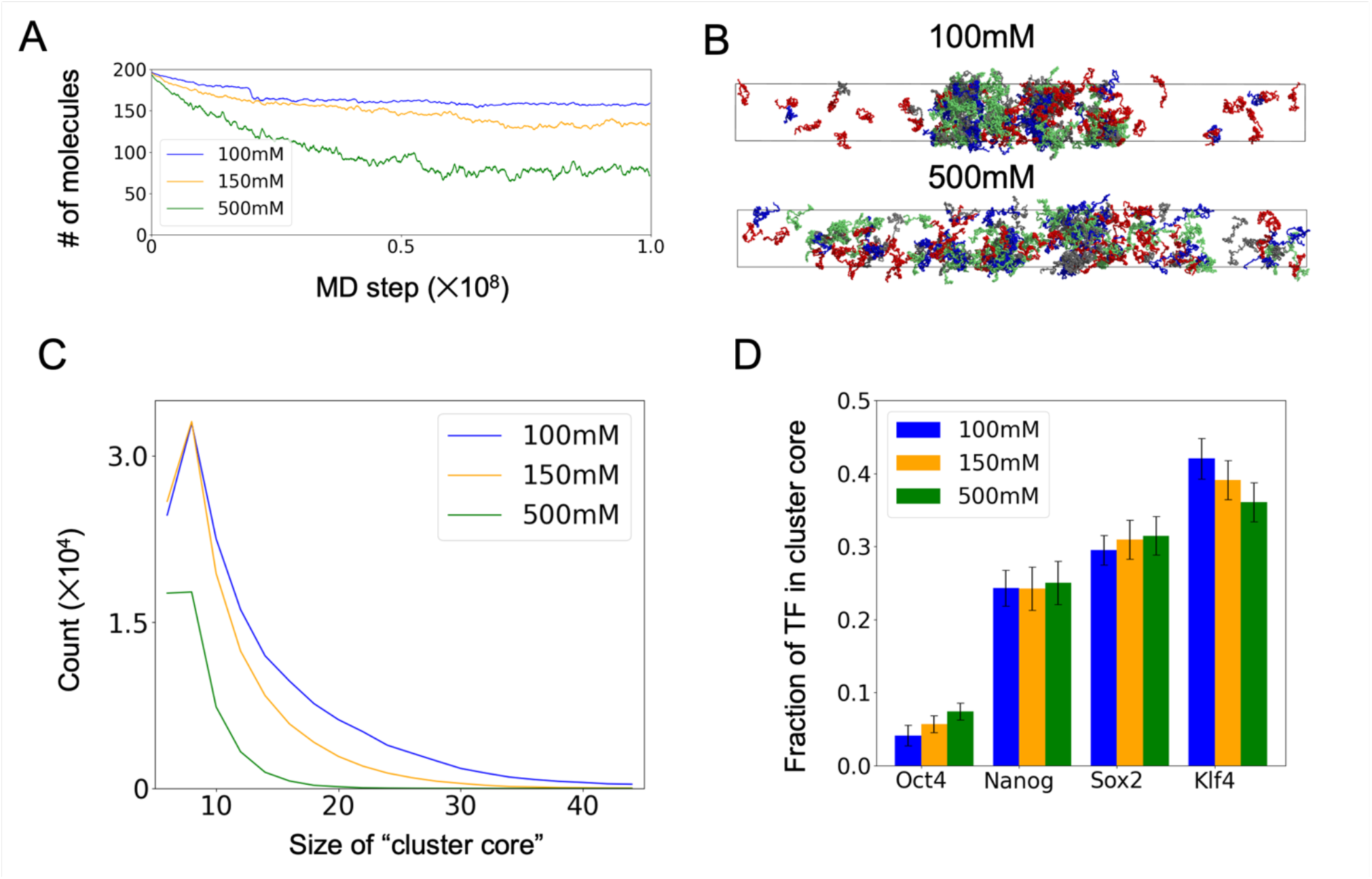
Salt-dependent simulations of the OSKN mixture. The results of the simulations with three salt concentrations, 100mM, 150mM, and 500mM, were compared (data for 150mM are the same as Fig.3). (A) The moving average of the number of molecules in the cluster. The results were averaged over five trajectories. (B) The snapshots at the end of each simulation with the salt concentration of 100 mM (top panel) and 500 mM (bottom panel). In each snapshot, Nanog, Oct4, Sox2, and Klf4 are colored with blue, red, gray, and green, respectively. (C) The histogram of the size of cluster cores. (D) The fraction of TFs in cluster cores. The error bars represent the standard errors.

To verify this hypothesis, we compared the number and size of the “cluster cores” in the three salt cases (Fig. 6C). We found that the number and the size of cluster cores decreased as the salt concentration increased. The result suggests that contrary to our prior expectation, electrostatic interactions play marked roles even in the formation of the cluster cores. TFs have negatively charged patches (around 100 residues in Klf4 and 250-275 residues in Nanog) and positively charged DNA-binding domains. Therefore, the cluster cores are stabilized both via hydrophobic and electrostatic interactions.

Next, we examined the salt-concentration dependence of fractions of TFs in the cluster core (Fig. 6D). The fraction of Klf4 decreased as the salt concentration increased, suggesting that the contribution of the electrostatic interactions is relatively strong for Klf4.

### Scaffolds and clients in OSKN condensate

In the simulations of the OSKN mixture, we found that the condensates were formed mainly by SKN, which coincides with the set of TFs that formed condensates in the simulations of individual systems. Then, how are the condensates formed by the mixture different from those formed in individual systems? While Oct4 did not form the condensate separately, was it recruited or excluded from the condensates formed by SKN? To examine these effects, we compared the amino acid densities along the Z-axis in the mixture and individual systems. We counted the number of amino acids in every bins of the size 300Å×300Å×60 Å along the Z-axis for the last 1000 frames of all trajectories and averaged them over the frames.

Figure 7A shows the amino acid densities of each TF ρ_aa_ along the Z-axis in the simulations of the OSKN mixture relative to the uniform density ρ_0_ of the corresponding TFs. As expected, SKN showed marked enrichment in the central 1000Å region, while Oct4 did not show large enrichment. As noted above, Oct4 was not excluded from the condensate either.

**Figure 7.**
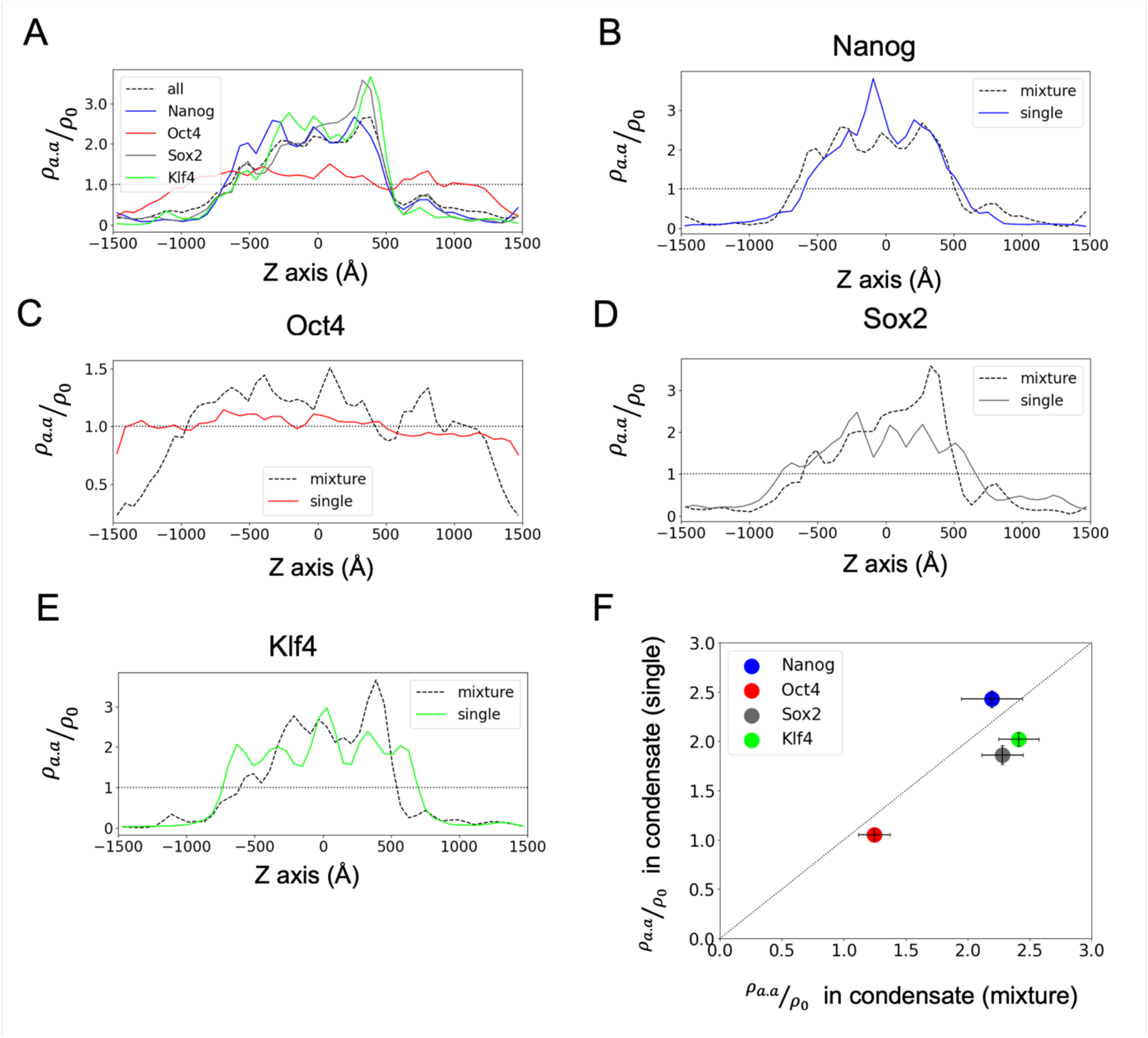
Comparison of the individual and OSKN mixture simulations. (A)The amino acid densities of a TF along the Z-axis (ρ_aa_) relative to those of uniform distribution (ρ_0_) in the OKSN mixture simulation. Colored solid curves represent the results of individual TFs, and the dashed black curve is the sum of all four TFs. (B-E) Comparison of the amino acid densities (ρ_aa_/ρ_0_) along the Z-axis of simulations between those with one TF and the OSKN mixture. (F) Comparison of the amino acid densities in the condensed phase (the Z coordinate of amino acids being between -500 and 500Å) for each TF in the individual systems (single) and the OSKN mixture (mixture). The error bars represent the standard errors between trajectories.

We found that in the cases of Klf4 and Sox2, the maximum densities were higher in the simulations of the mixture than in the individual simulations, while Nanog showed an opposite result (Fig. 7B, D, E). For Klf4 and Sox2, the electrostatic interaction plays key roles in the condensates where interactions between hetero molecules are favorable. Especially, Sox2 itself has only a modest tendency to form condensates but showed enrichment in the condensates in the mixture system via primary interaction to Klf4 (see also Fig.4B). On the other hand, as shown in our previous study, Nanog in the individual system forms micelle-like clusters primarily via the hydrophobic interactions in its WR regions. Thus, Nanog could form more compact structures by binding to other Nanog molecules at the center of the clusters. Thus, the maximum density became higher in the simulations with Nanog alone.

Figure 7C showed that the Oct4 density was slightly enriched within and around the condensates of the mixture mainly formed by SKN. This result suggests that Oct4 can be recruited, albeit moderately, by SKN. Referring to Fig.4B, the recruitment can mainly be made by Sox2 and Klf4. In Fig.7C, Oct4 density drops only near ± 1500Å, making the interpretation subtle. To confirm the moderate enrichment of Oct4 around the condensate region, we expanded the Z-side length of the simulation box to 5000 Å and extended the simulation starting from the final configuration of the above simulation (Fig. S10). The resulting amino acid densities of each TF ρ_aa_ are plotted in Fig.S10E, which confirmed a moderate enrichment of Oct4 density around the condensates formed by SKN.

To better quantify the comparison, we calculated the fraction of molecules in the condensates defined simply as -500Å < Z < +500Å in the mixture and individual systems (Fig. 7F). Results suggest that Sox2 and Oct4 were enriched in the mixture system relative to the individual systems. The density of Klf4 also increased when mixed with other molecules, which suggested that the mixture of TFs not only recruited other molecules but also increased the density of the components. On the other hand, Nanog showed nearly the same tendency of condensates between the mixture and individual systems. Based on these results and those in Fig. 4B, we suggest that Klf4 worked as the scaffold of the condensate, while Sox2 was enriched by hetero-molecular interactions with Klf4, and Oct4 was weakly recruited to the condensate.

## DISCUSSION

In this study, we investigated heterogeneous organization and interactions in condensates formed by four TFs, Nanog, Oct4, Sox2, and Klf4, which have crucial roles in ESCs using residue-resolution coarse-grained model simulations. In the individual systems, Nanog, Sox2, and Klf4 separately formed stable clusters, while Oct4 molecules diffused to the entire slab box. In the mixture system, the three TFs, Sox2, Klf4, and Nanog, formed cluster cores mainly by the interactions between their relatively hydrophobic IDRs. The relatively hydrophobic IDRs were in the center of the clusters, while DNA-binding domains and hydrophilic IDRs were on the surface or outside of the clusters. This feature is like the clusters formed by Nanog alone, which we found in the previous study(16). Comparing the simulations for the mixture of four TFs and for individual systems, we suggest that Klf4 served as the scaffold, while Sox2 was enriched by its interaction with Klf4, and Oct4 was moderately recruited via its interaction mainly with Sox2 and Klf4.

What can be the functional roles of such a heterogeneous organization of TFs? In the transcriptional condensation model, the condensates formed by TFs, together with Mediators, coactivators, and RNA polymerases, provide a micro-environment to communicate between multiple enhancers and promotors over long distances. Heterogeneous organization in condensates can provide locally distinct multiple local environments, each enriching different molecules for different functions. For example, condensates used for transcription initiation and those for transcription elongation can be distinct, facilitating the transition in the transcription cycle(23). Phosphorylation of TFs and other proteins can control the reorganization of the condensates. Obviously, there is a long way to obtain a complete picture of such controls.

In this study, we focused only on the interactions of four TFs, but there are also interactions with other TFs or other transcription-related proteins in the transcriptional condensate in the cell nuclei. Mediator is an important factor in the condensate, and Boija et al. showed that Mediator could also show LLPS and recruit other transcription-related proteins, including those used in this study(5). The C-terminal tail of RNA Polymerase II is also known to form droplets from *in vitro* experiments, and the phosphorylation of Serine or Threonine residues in the tail regulates the transcription activity(24). These proteins can affect the configurations and dynamics of the condensates formed by TFs. The effects of these molecules on the condensate are of interest to future studies.

## METHOD

### Molecules

We used the full-length sequences of four mouse TFs, Oct4, Sox2, Klf4, and Nanog.

### Coarse-grained Models for Transcription Factors

We used the residue-resolution coarse-grained model for large-scale MD simulations of TF condensates. For each TF, each amino acid was represented by one bead located at the Cα atom position. Since each TF has one DNA-binding domain and IDRs, we used the force-field AICG2+ for DNA-binding domains and the flexible local potential for disordered domains to represent the local conformations(25, 26). The AICG2+ model was used to maintain the folded structures. We employed Modeller(27) to generate all-atom structures from PDB (2VI6(28) for Nanog and 4M9E(29) for Klf4) and used them as the reference structures of Nanog and Klf4. In the case of Sox2 and Oct4, we used the structures predicted by AlphaFold2 as the reference(30). For generic inter- and intra-molecular interactions, we used the excluded volume interactions, electrostatic interactions, and the hydrophobicity scale (HPS) model(11). The HPS model was applied to the interactions between IDRs of intra- and inter-molecules. Oct4 has a short linker region between two DNA-binding domains, which was treated as a part of the globular domain, without using the HPS potential. Among several parameter sets of the HPS model(13, 14, 31), we chose the parameter set obtained by Tesei et al.(14).

### Molecular Dynamics Simulation

As in previous simulation studies of LLPS, we used a long rectangular simulation box (small X and Y side lengths and a large Z side length) with the periodic boundary condition. This rectangular box is superior to the cubic box for the phase-separated systems in that the slab configuration reduces the finite-size effect only to the interface normal to the Z-axis. With the standard cubic box, the droplet, once formed, has a spherical surface. We used a slab box with X, Y, and Z side lengths of 300, 300, and 3000 Å, respectively. From a preparatory simulation of one TF molecule, we estimated the largest residue-residue distance within one molecule as ∼250 Å. Thus, we used 300 Å as the X and Y side lengths to prevent the molecules from interacting with their periodic images. To confirm it in the mixture simulation of the mixture, we calculated the largest residue-residue distance within every molecule. The distance distribution had a peak at 150 Å and the largest value at 250 Å, ensuring that 300Å box size in the X and Y sides are large enough. As for the Z side length, we avoid too large side to prevent intractable equilibrium. We checked single molecules can diffuse ∼ 3000 Å within the possible simulation time. For the simulations of individual TF, we included 200 molecules of one TF. For the OSKN mixture simulations, we included 50 molecules of each TF. The total protein density in both systems corresponds to an average concentration of 1.23 mM. This average concentration is selected to between the densities of the dense and dilute phases of the molecules, ensuring phase separation. We used the MD simulation software GENESIS ver2.1.0(32-34), which contains all the models described above and used GENESIS-CG-TOOL(34) to generate CG MD input files. All simulations in this study used Langevin dynamics at a temperature of 300 K.

In both the individual and mixture simulations, we prepared completely phase-separated configurations as starting configurations of the product runs. First, we prepared the configuration in which 200 molecules were located on grids, each of which was in a compact conformation, avoiding any overlaps. Then, we conducted short MD runs with the box size gradually decreasing by 0.1Å every 100 MD steps along the z-axis, until the size was small enough for all molecules to be in the condensates. In this shrinking simulation, we used the same force field as the one used in the product runs. For the OSKN mixture simulations, we performed additional simulations to relax the condensate configurations with the Z side lengths of 600 Å, which is large enough for TFs to diffuse in the simulation box.

As product runs, we performed five independent MD runs for monovalent salt concentrations of 150, 100, and 500 mM, for the OSKN mixture and for the single-TF systems.

### Analysis

In most analyses, we used Python-library MDAnalysis(35, 36) to read DCD files and analyze trajectories. To quantitatively evaluate the size of clusters, we defined them based on the number of intermolecular contacts. We first calculated the residue-residue contact map for each pair of molecules with a residue-residue cutoff distance of 6.5 Å and counted the number of contacts in every frame. We defined the contacts between two molecules under the condition that the number of residue-residue contacts was larger than three to reduce the accounting of chance-based transient contacts. We considered molecules being in the same cluster when they are in contact. Then, we defined a cluster containing more than 20 molecules as a condensate.

We defined the “cluster core” with a more stringent criterion to find a robust central part of clusters. We used 10, rather than three, as the cutoff number of residue-residue contacts for the “cluster core”. After clustering, we selected the cluster cores that contained five or more molecules in the cluster core, which were used for later analysis.

In the FRAP-like analysis(19, 20), we selected molecules in the same bin at a frame and calculated the standard deviations of the Z-coordinate of the selected molecules’ center of mass over the next 1000 frames. The mean square deviation increased linearly over time, suggesting the normal, not the anomalous, diffusion of proteins.

## AUTHOR CONTRIBUTIONS

The manuscript was written through contributions of all authors.

## DECLARATION OF INTERESTS

The authors have declared that no competing interests exist.

## ACKNOWLEDGEMENT

This work used computational resources of supercomputer Fugaku provided by the RIKEN Center for Computational Science (Project ID: hp230209, hp240215). Giovanni B. Brandani originally created the CG MD input files for Klf4.

## Notes

### Competing Interest Statement

The authors have declared no competing interest.

### Summary of Updates

We have reformatted the paper to make it easier to read and revised some figures and references.

