## Supplemental Figures for "Heterogenous organization in condensates of multiple transcription factors in embryonic stem cells"

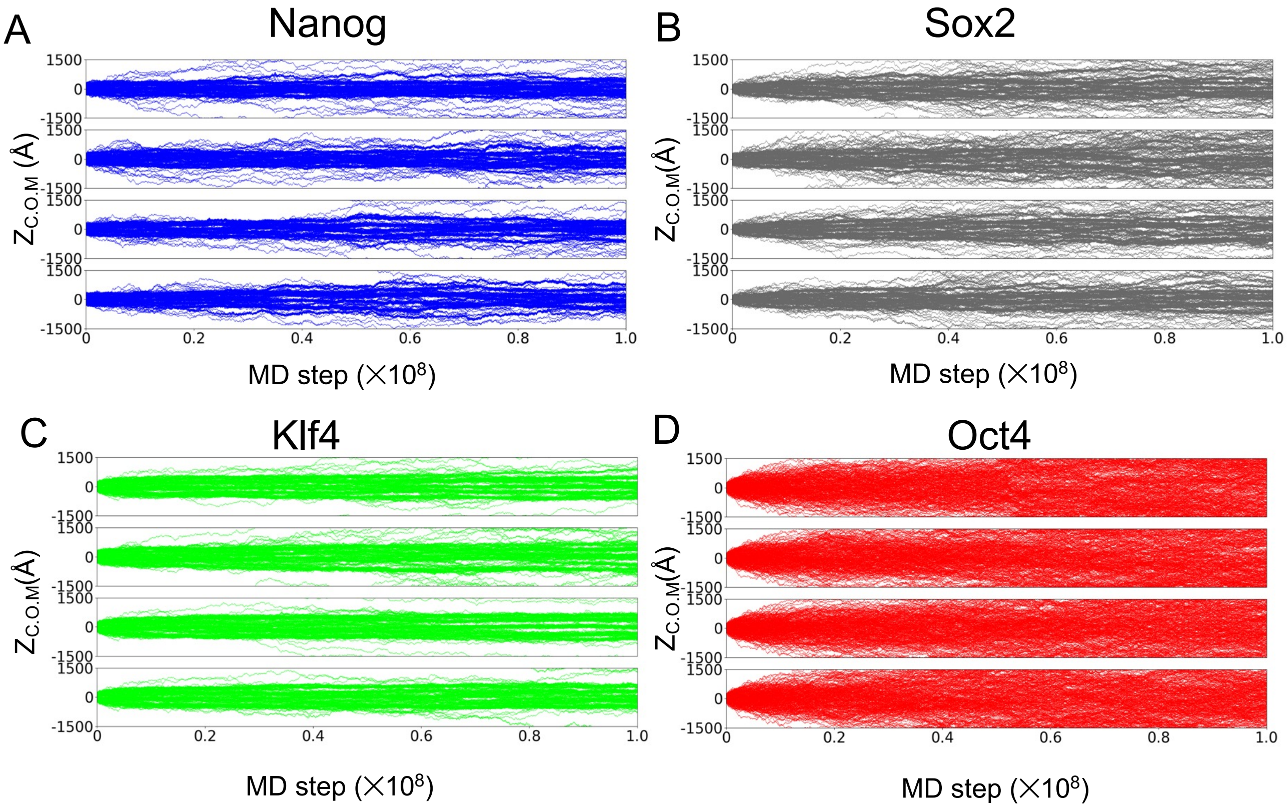


Figure S1. The time courses of Z-coordinates of the center of mass of individual molecules in individual systems (A: Nanog, B: Oct4, C: Sox2, D: Klf4). For each of TF, four trajectories not shown in the top panels of Fig. 2 were plotted.

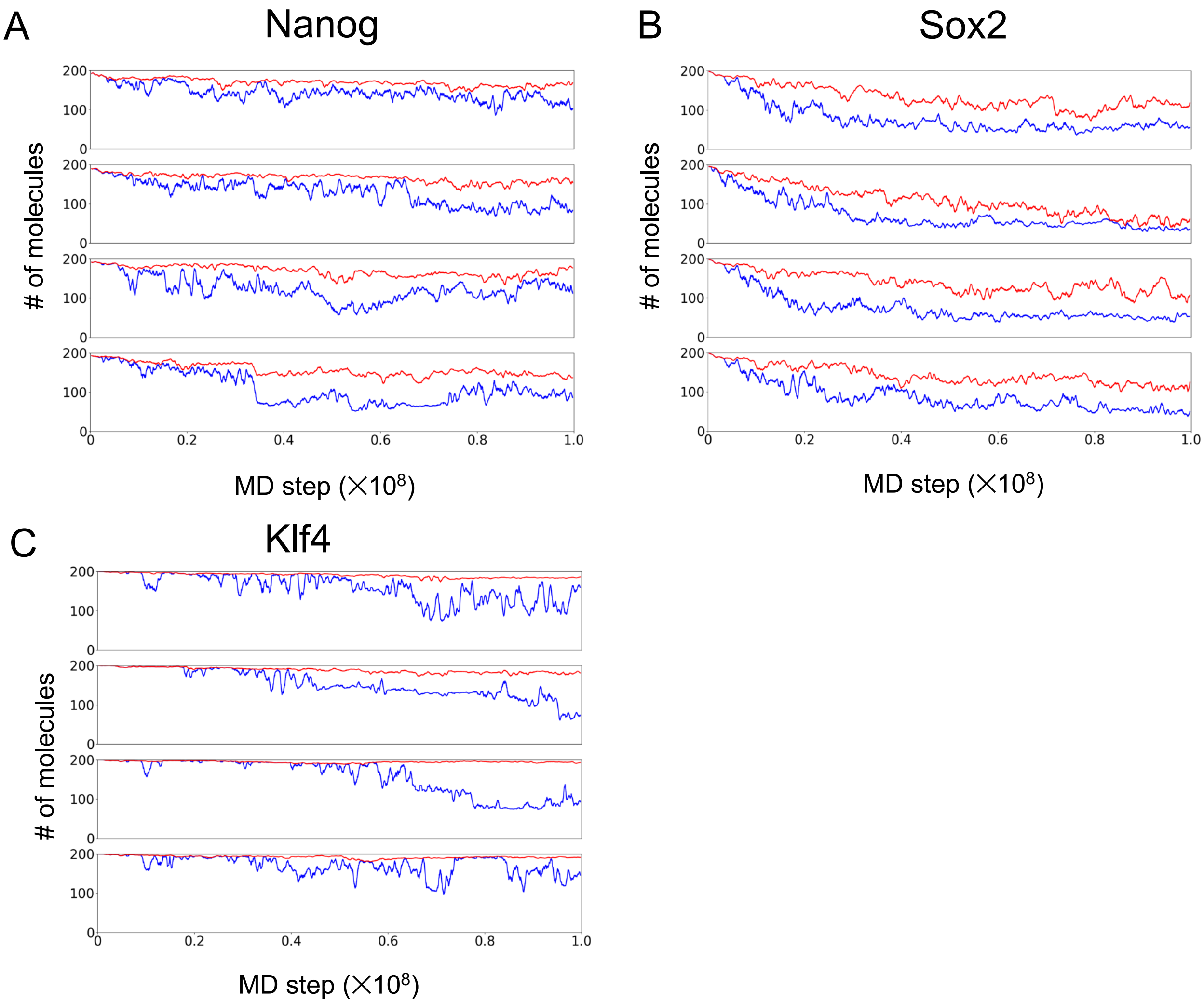


Figure S2. The moving-average of molecules for each TF in the clusters (red) and those in the largest cluster (blue) in individual systems. For each of TF, four trajectories not shown in the middle panels of Fig. 2 were plotted. This analysis was not performed for Oct4 because it did not form droplets.

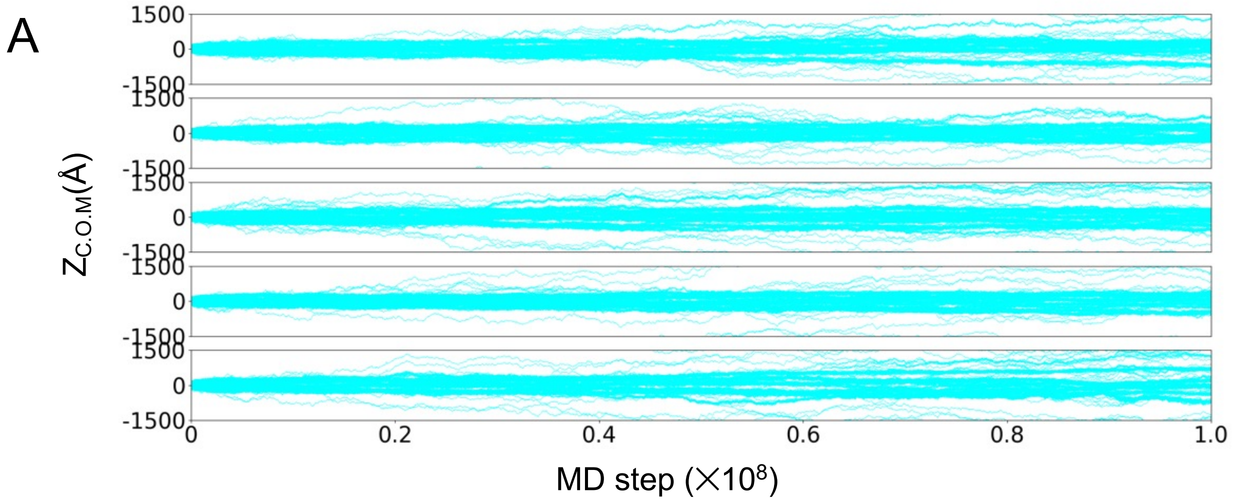


Figure S3. Simulations of the system containing 200 molecules of human Nanog under conditions of 150mM NaCl concentration. To compare with our previous study about human Nanog, we performed the LLPS simulations of human Nanog under the same conditions as this study. (A) The time courses of Z-coordinates of the center of mass of individual molecules. Each panel represents the result from the trajectory with different stochastic force

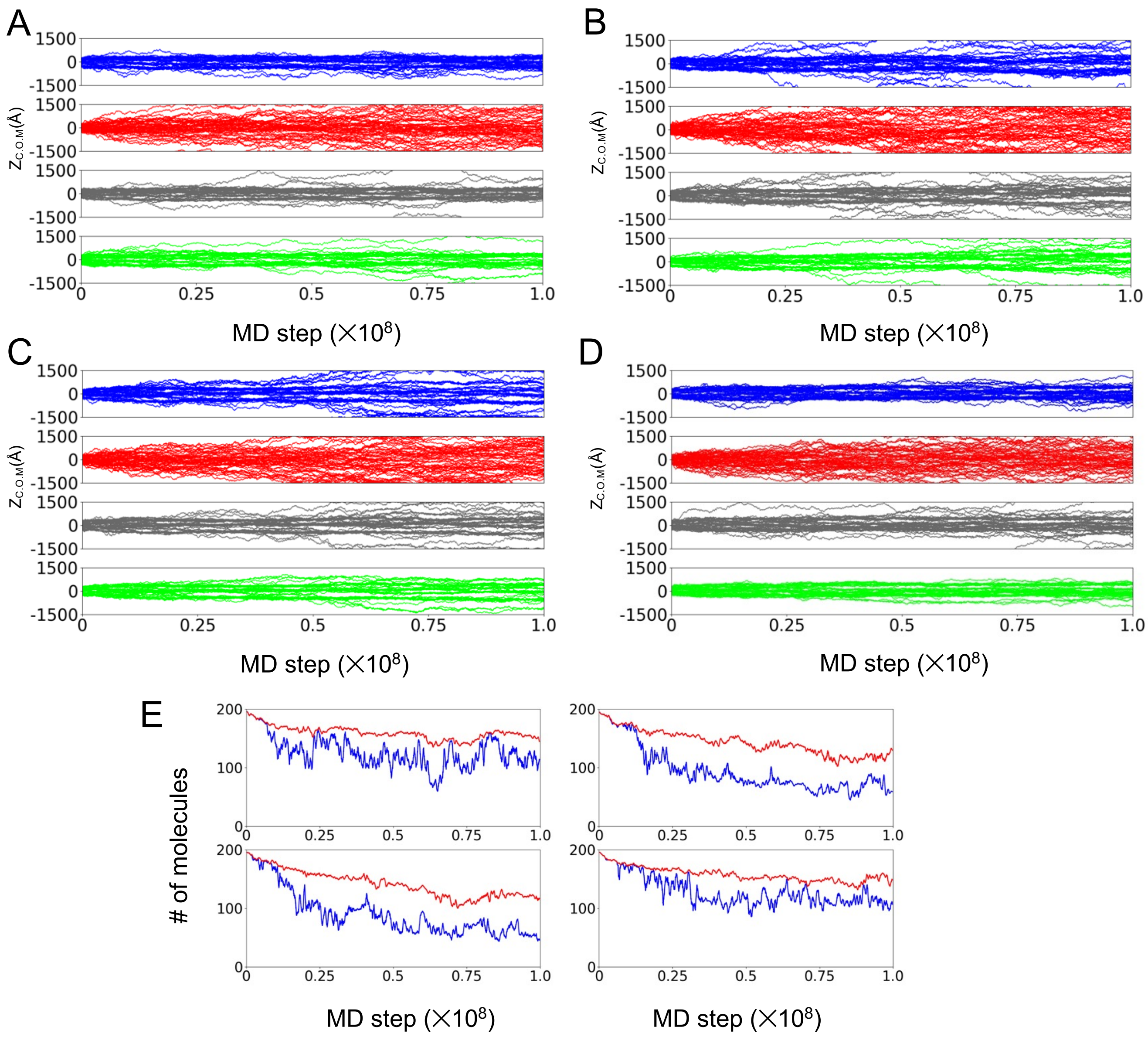


Figure S4. Simulations of OSKN mixture containing 50 molecules of each TF. Four trajectories not shown in Figure 3 are plotted. (A~D) The time courses of Z-coordinates of the center of mass of individual molecules. Each panel shows the result of Nanog, Oct4, Sox2, and Klf4 from the top to the bottom. (E) The moving-averaged numbers of molecules in the clusters (red) and those in the largest cluster (blue). Each panel shows the results obtained from the same simulation as the data in A~D.

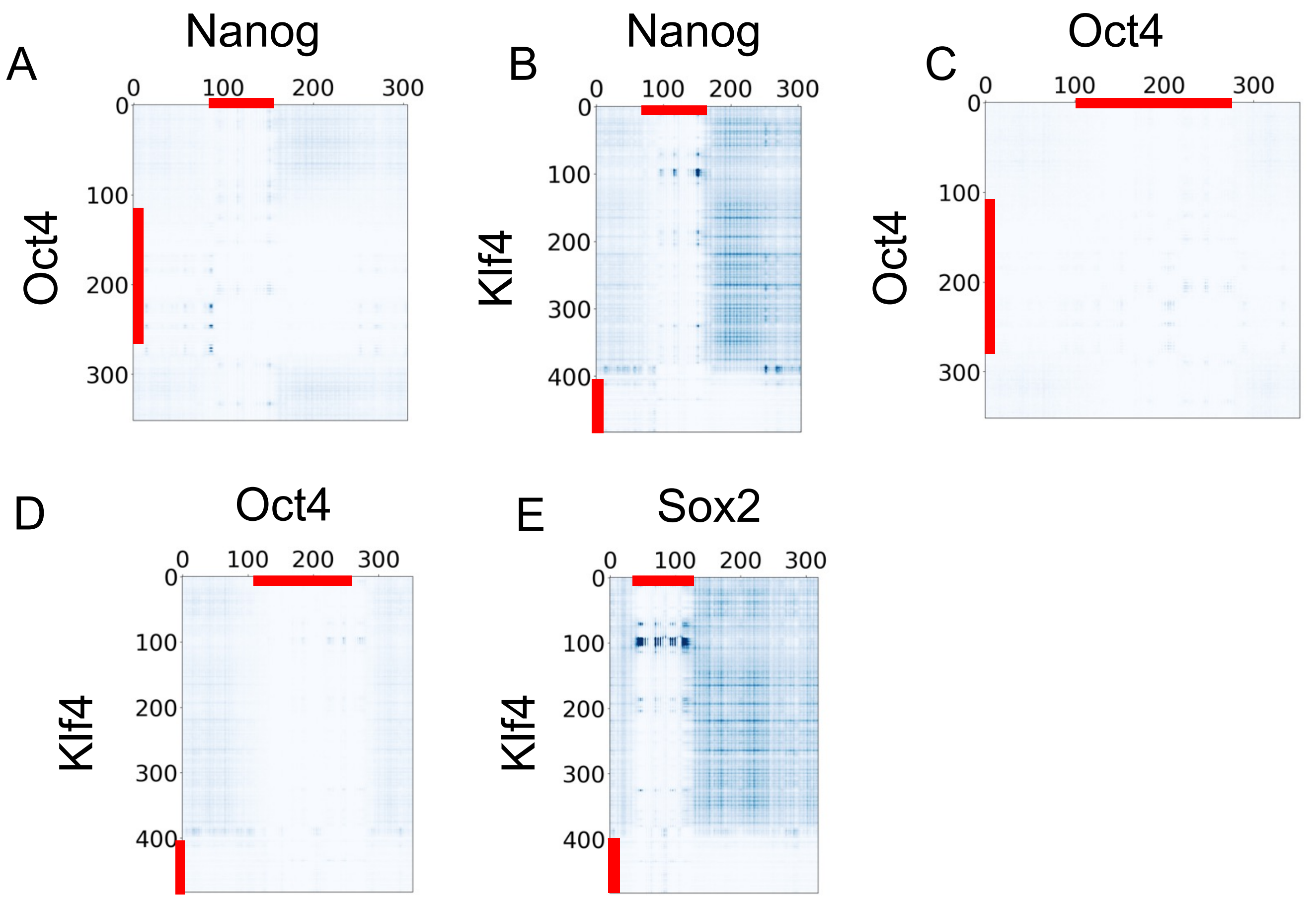


Figure S5. Residue-wise TF-TF interactions not shown in Figure 5A-C. (A~E) The average number of contacts between residues of each pair of TFs. The red lines represent the DNA-binding domains.

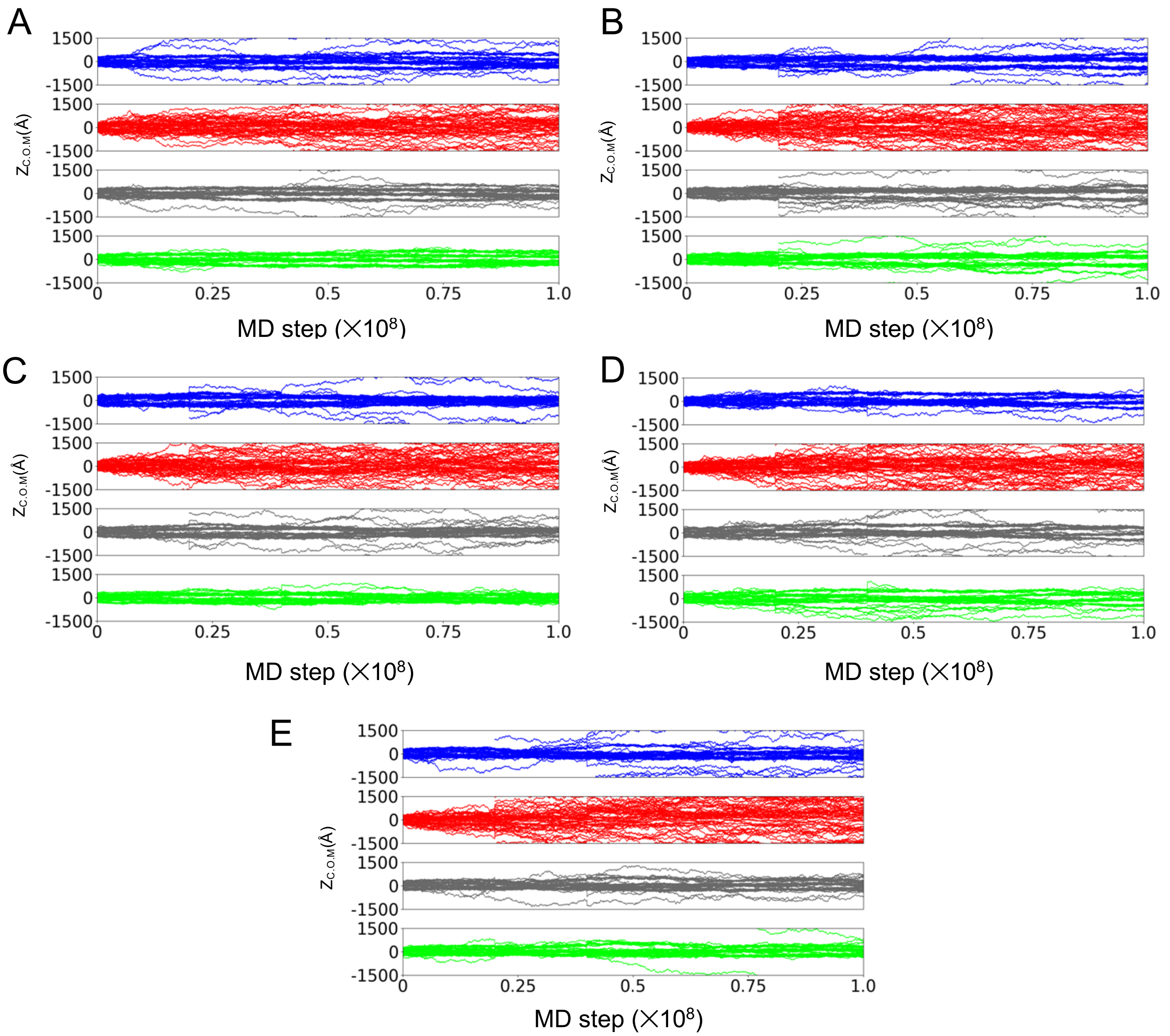


Figure S6. The time courses of Z-coordinates of the center of mass of individual molecules under the condition with 100 mM salt concentration. In A~D, each panel shows the result of Nanog, Oct4, Sox2, and Klf4 from the top to the bottom. Figure A~D represent the results from the trajectories with different stochastic forces.

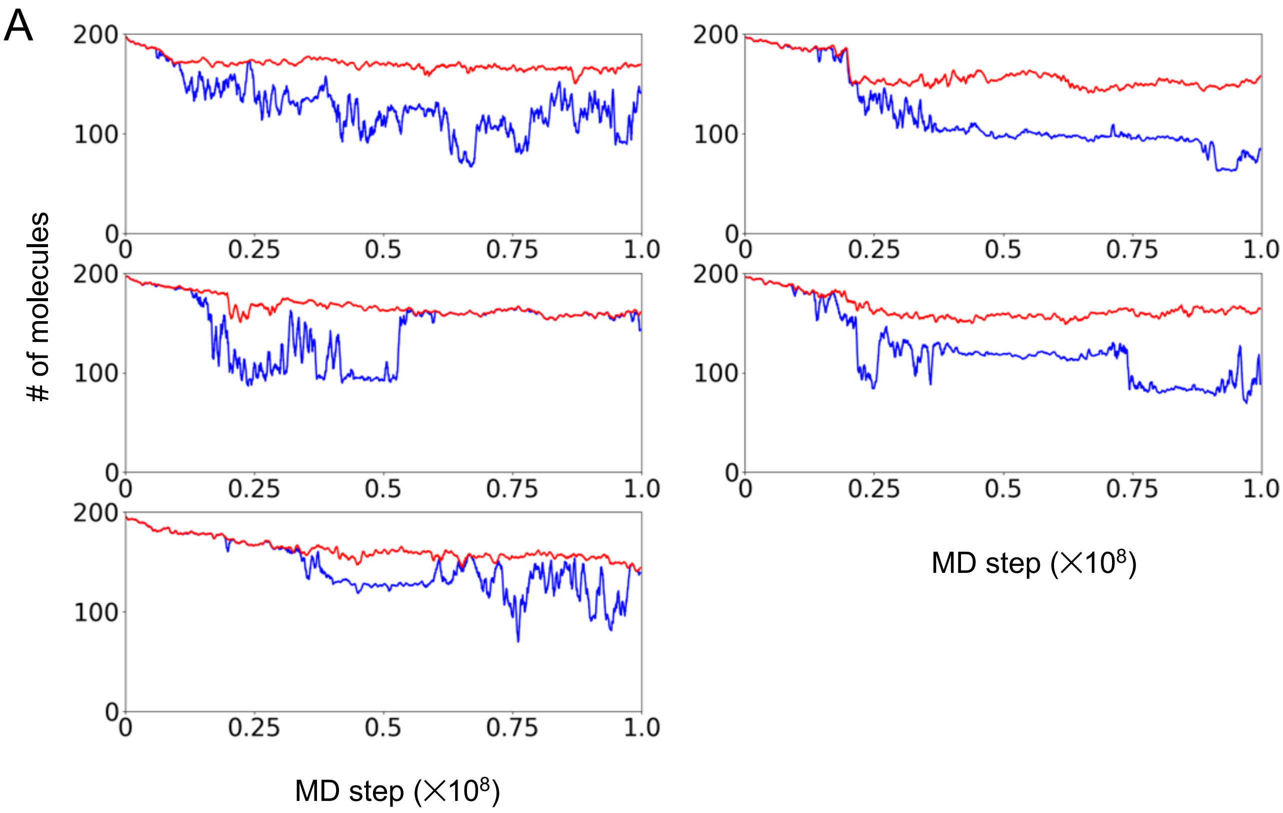


Figure S7. The moving-average of the number of TF molecules in the clusters (red) and those in the largest cluster (blue) under the condition with 100 mM salt concentration. The results of this figure are based on the same simulation data as Fig. S6.

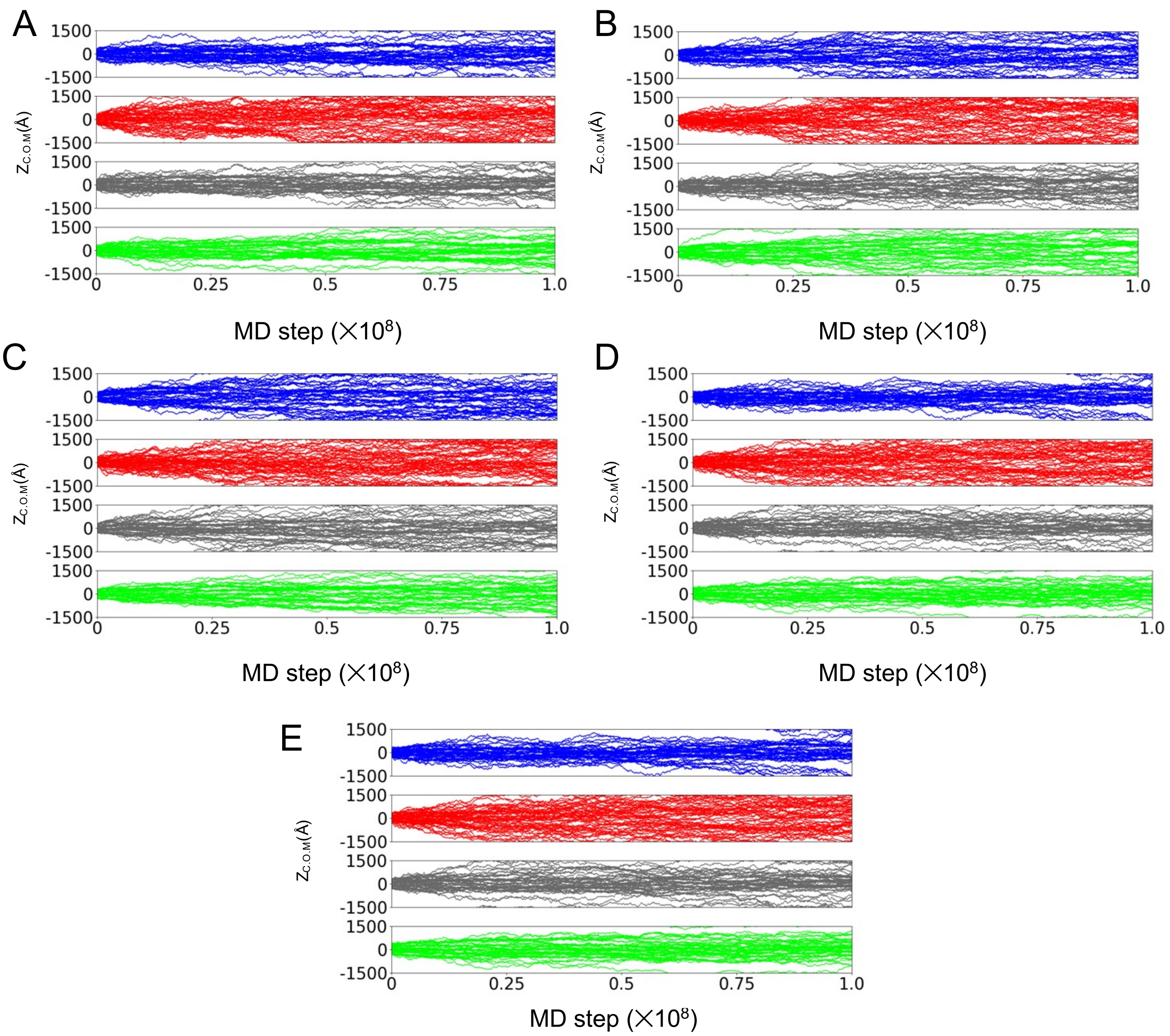


Figure S8. The same as Figure S6 with 500 mM.

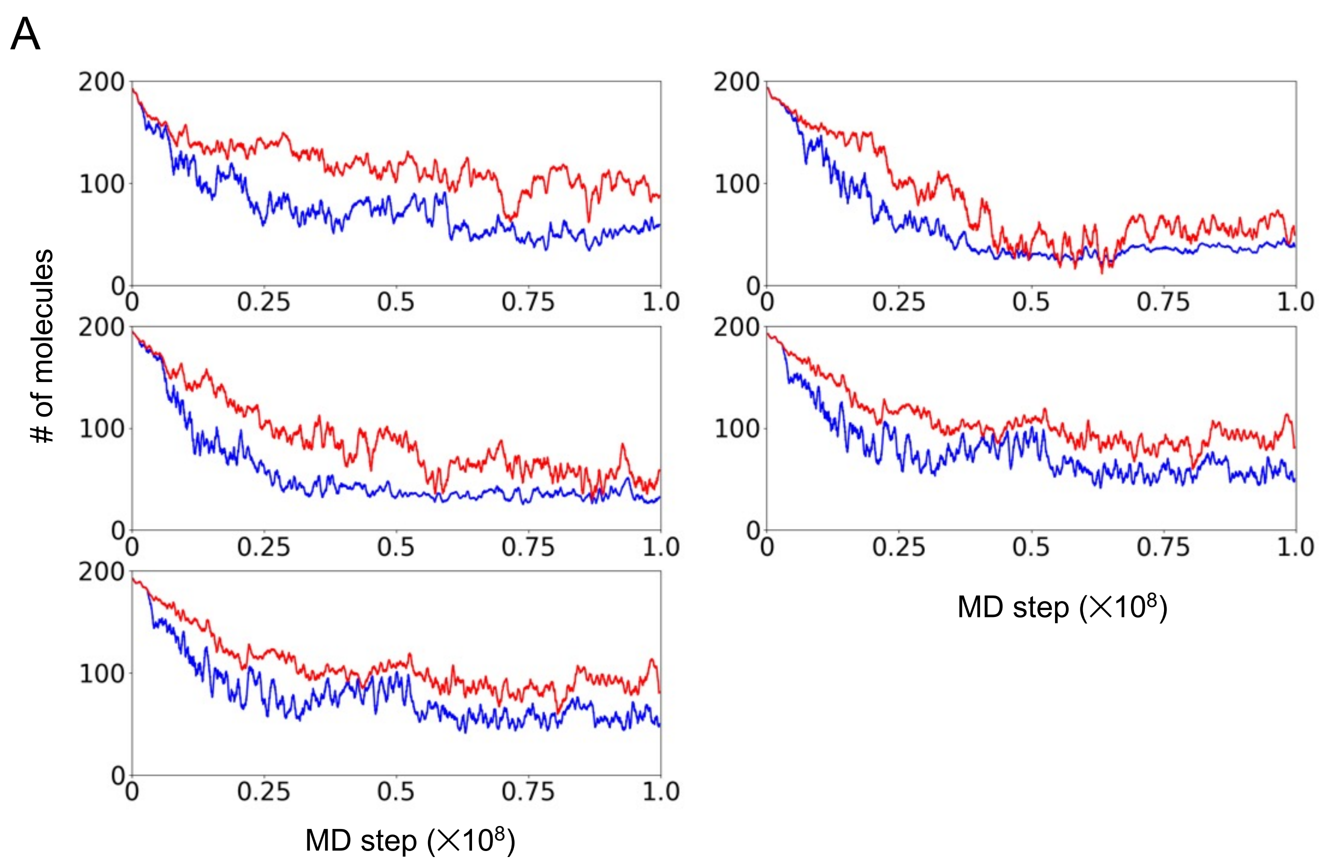


Figure S9. The same as Figure S7 with 500 mM.

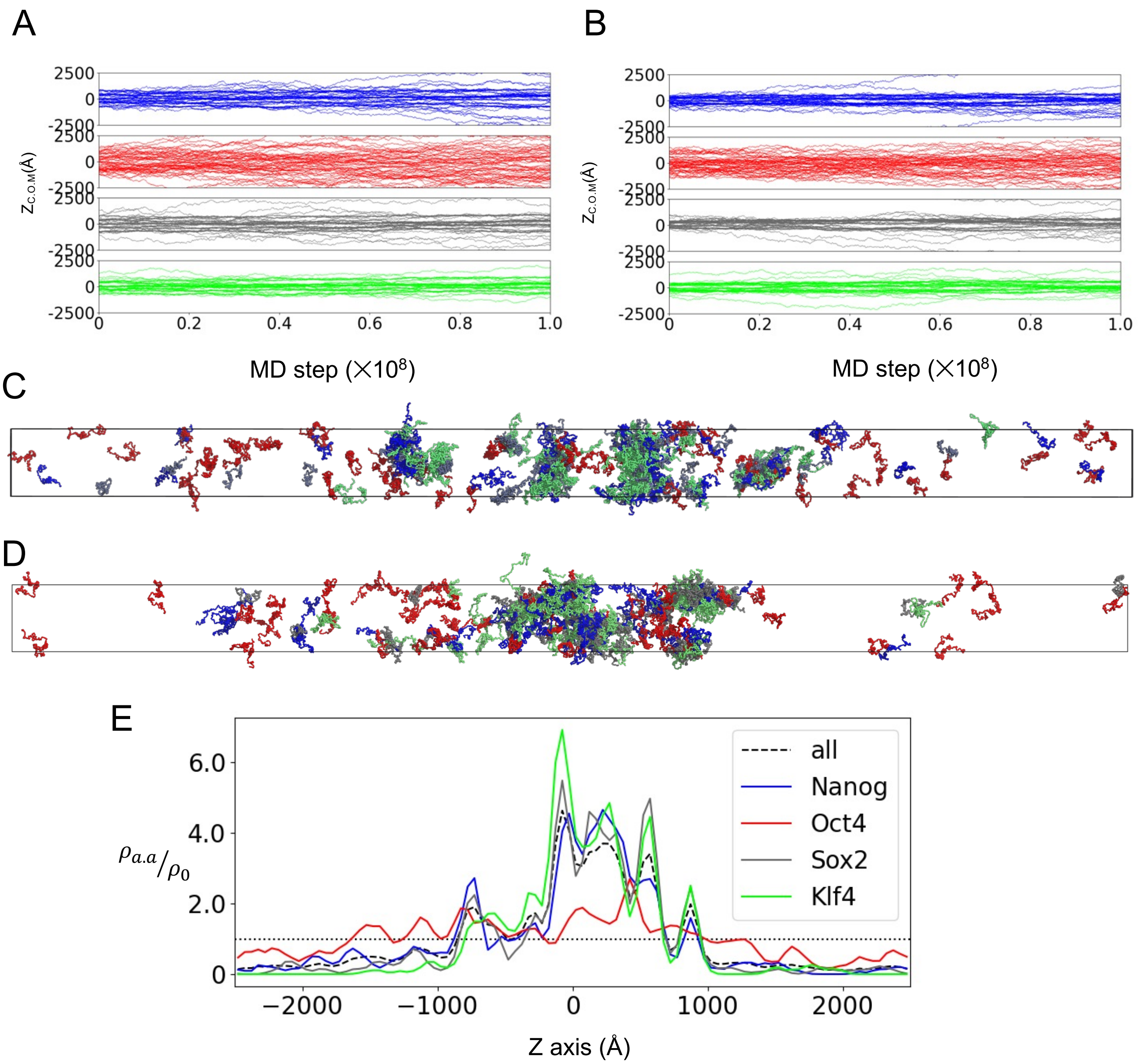


Figure S10. Simulations of OSKN mixture containing 50 molecules of each TF with an elongated box size (300Å×300Å×5000Å). In the simulations, we used the final structures performed under the conditions with 150mM NaCl concentration as the initial structures. (A, B) The time courses of Z-coordinates of the center of mass of individual molecules. In the figures, each panel shows the result of Nanog, Oct4, Sox2, and Klf4 from the top to the bottom. (C, D) The snapshots of the final structures in the same trajectory as those in Fig. S10A. In each snapshot, Nanog, Oct4, Sox2, and Klf4 were colored blue, red, gray, and green. (E)The amino acid densities of a TF along the Z-axis ($\rho_{aa}$) relative to those of uniform distribution ($\rho_0$) in the OKSN mixture simulation. Colored solid curves represent the results of individual TFs, and the dashed black curve is the sum of all four TFs.
